# *Salmonella* Typhimurium Manipulates Syntaxin 7 to Navigate Endo-Lysosomal Trafficking in Host Cells

**DOI:** 10.1101/2024.12.12.628128

**Authors:** Rhea Vij, Ritika Chatterjee, Abhilash Vijay Nair, Anmol Singh, Dipasree Hajra, Subba Rao Gangi Setty, Dipshikha Chakravortty

**Affiliations:** Department of Microbiology and Cell Biology, Division of Biological Sciences, Indian Institute of Science, Bangalore, Karnataka, India; Adjunct Faculty, Indian Institute of Science Research and Education, Thiruvananthapuram, Kerala, India; Present address: Laboratory of Biomedical Mass Spectrometry, Biosciences Institute, Newcastle University, Newcastle upon Tyne, UK; Present address: Department of Microbiology and Immunology, Vagelos College of Physicians and Surgeons, Columbia University, New York, NY 10032, USA

**Keywords:** *Salmonella*-containing vacuoles, macrophages, epithelial cell, syntaxin 7, SNAREs

## Abstract

Intracellular pathogens rely on manipulating host endocytic pathways to ensure survival. *Legionella* and *Chlamydia* exploit host SNARE proteins, with *Legionella* cleaving syntaxin 17 (STX17) and *Chlamydia* interacting with VAMP8 and VAMP7. Similarly, *Salmonella* targets the host’s endosomal fusion machinery, using SPI effectors like SipC and SipA to interact with syntaxin 6 (STX6) and syntaxin 8 (STX8), respectively, maintaining its vacuolar niche. Recent evidence highlights syntaxin 7 (STX7), a Qa-SNARE involved in endo-lysosomal fusion, as a potential *Salmonella* target. BioID screening revealed STX7 interactions with SPI-2 effectors SifA and SopD2, suggesting a critical role in *Salmonella* pathogenesis. We investigated the role of STX7 in *Salmonella*-containing vacuole (SCV) biogenesis and pathogenesis in macrophages and epithelial cells. Our findings indicate that STX7 levels and localization differ between these cell types during infection, reflecting the distinct survival strategies of *Salmonella*. Live cell imaging showed that STX7 is recruited to SCVs at different infection stages, with significantly altered distribution in HeLa cells at the late stage of infection. STX7 knockdown resulted in reduced bacterial survival, which was rescued upon overexpression of STX7 in both HeLa and RAW264.7 cells, suggesting *Salmonella* hijacks STX7 to evade lysosomal fusion and secure nutrients for intracellular replication. These results underscore the essential role of STX7 in maintaining SCVs and facilitating *Salmonella* survival. Further, the temporal expression of STX7 adaptor/binding partners in macrophages showed dynamic interactions with STX7 facilitating *Salmonella* infection and survival in host cells. Together, our study highlights STX7 as a critical host factor exploited by *Salmonella*, providing insights into the molecular mechanisms underlying its pathogenesis in macrophages and epithelial cells. These findings may in form strategies for targeting host-pathogen interactions to combat *Salmonella* infections.

## 1. Introduction

Many recent studies emphasize the importance of manipulating host endocytic pathways for the survival of intracellular pathogens [1]. It has been reported that *Legionella* and *Chlamydia* modulate host SNAREs to establish infection and ensure their survival. The *Legionella* effector Lpg1137 cleaves syntaxin 17 (STX17), while the IncA protein of *Chlamydia* contains SNARE-like motifs and interacts with vacuole-associated membrane protein 8 (VAMP8) and vacuole-associated membrane protein 7 (VAMP7) [2, 3]. *Salmonella* exploits the host endosomal fusion machinery to access host membranes and nutrients essential for its intracellular survival and replication [4]. Specifically, *Salmonella* SPI effectors SipC and SipA interact with syntaxin 6 (STX6) and syntaxin 8 (STX8) to maintain their vacuolar niche, enabling their survival within the host [5, 6]. Recent studies from our lab also demonstrated that *Salmonella* hijacks syntaxin 3 (STX3) via SPI-2 effectors and alters the expression of STX17 to mediate its survival in the host cell [7, 8]. Additionally, *Salmonella* regulates vacuole size by fusing with SNAP25- positive infection-associated macropinosomes in a syntaxin 4-dependent manner [9].

An *in silico* analysis predicted interactions between *Salmonella* effectors and host syntaxins, including syntaxin 7 (STX7) [10], a SNARE involved in endo-lysosomal fusion [11]. This was validated through a BioID screen of *Salmonella* T3SS-2 effectors, which revealed that STX7 interacts with SifA and SopD2 [12]. Consequently, we aim to investigate the role of STX7 in the intracellular pathogenesis of *Salmonella*. STX7 belongs to the Qa-SNARE subfamily and is primarily located on late endosomes and lysosomes [11]. Some studies also report its presence on the plasma membrane, early endosomes, and recycling endosomes [13, 14]. STX7 forms a complex with Vti1b (Qb), Stx8 (Qc), and VAMP8 (R), which mediates homotypic late endosome fusion. When VAMP7 replaces VAMP8 in this complex, it mediates the fusion of late endosomes with lysosomes [15]. STX7 also facilitates phagosome-lysosome fusion [16] and mediates vesicular trafficking from the trans-Golgi network to endosomes [17]. Its activity is regulated by post-translational modifications, such as phosphorylation induced by colony-stimulating factor-1, which enhances the formation of the STX7-Vti1b-STX8-VAMP8 SNARE complex in macrophages [18]. STX7 interacts with the HOPS and CORVET tethering complexes and their positive UVRAG (UV radiation resistance-associated gene) regulator. Influenza A virus exploits this interaction to assemble the STX7-Vti1b-STX8 SNARE complex and recruit VAMP8-positive endosomes, facilitating viral entry and survival [19]. Similarly, *Helicobacter pylori*, an intracellular pathogen, targets STX7-mediated trafficking pathways, residing within a hybrid late endosomal-lysosomal vacuole that supports bacterial survival [20]. *Salmonella*, on the other hand, modulates its vacuole to prevent lysosomal fusion. STX7 knockdown has been shown to impair the formation of SIFs (*Salmonella*-induced filaments) in *Salmonella*-infected HeLa cells [21]. A recent report by Pha et al. demonstrates that IncE (an inclusion protein of *Chlamydia trachomatis*) interacts with STX7 and STX12 through its SNARE-like domain to access host cargos [22]. Therefore, we aim to investigate the role of STX7 in SCV biogenesis and *Salmonella* pathogenesis. We report that the levels of STX7 change during *Salmonella* infection in both macrophages and epithelial cells. However, not in a similar fashion; this could be potentially due to the differential survival ability of *Salmonella* in epithelial cells and macrophages. We also have shown here that the STX7 gets recruited onto the SCV at different stages of infections, and there is a significant difference between the cellular localization of STX7 upon infection in epithelial cells, e.g., HeLa cells. We also have further knocked down the STX7 from the cells, and we observed that there is reduced survival, suggesting that potentially *Salmonella* hijacks STX7 for its own benefit, such as avoiding lysosome while acquiring essential nutrients for its survival and replication. Together, our data strongly demonstrate the potential role of STX7 in the *Salmonella* pathogenesis and infection in both macrophages and epithelial cells.

## 2. Results

### 2.1 The level of STX7 shows bi-modal regulation during infection with STM WT in macrophages

Upon infection with wild-type *Salmonella* Typhimurium strain 14028S (STM WT) in RAW 264.7 cells, we observed that there is an increase in the transcript level of STX7 at the initial stage of infection (2h post-infection) (Fig.1A). During the intermediate phase of infection (6h post-infection), there is a decrease in STX7 transcript level (Fig.1A). At the late stage of infection (10h post-infection), there was a rise in the transcript levels of STX7 mRNA (Fig.1A). Further, we observed an increment in mRNA transcript levels of STX7 at 2h post-infection with PFA fixed bacteria compared to the uninfected control (Fig.1A), but this is significantly decreased in comparison to infection with the wild-type bacteria (Fig.1A). This can be attributed to the role of STX7 in mediating phagosome-lysosome fusion [16]. Additionally, we have performed immunofluorescence studies using confocal laser scanning microscopy (CSLM) in RAW 264.7 macrophages upon *Salmonella* infection to extend the findings to the protein level. We observed that the expression pattern exhibited by STX7 is similar to that of the transcript levels, i.e., expression increases 2h post-infection, followed by a decline at 6h and again a rise at 10h post-infection (Fig.1B, C). Thus, STX7 exhibits a bimodal expression profile at both the transcript and protein levels during *Salmonella* infection. *Salmonella* within the host cell resides in a modified phagosome known as the *Salmonella*-containing vacuole (SCV). The mature SCV bears multiple markers of the late endosome, e.g., LAMP1, vATPase, etc. [23]. Along the same line, several studies report that STX7 is present on late endosomes [24], suggesting that there might be a strong interplay of host STX7 during *Salmonella* infection. In HeLa cells, STX7 is recruited on the SCV 90 min post-infection with *Salmonella* [6]. We previously demonstrated that STX7 is upregulated at transcript and protein levels during *Salmonella* infection in RAW264.7 macrophages. Hence, we wanted to determine whether STX7 localizes on the SCV in macrophages using CSLM. This would also confirm the vacuolar status of the bacterium; LAMP1 acts as a marker for the SCV [25, 26]. We observed that the bacteria co-localizes with STX7 at the early, intermediate, and late stages of infection, i.e., at 2h, 6h, and 10h in RAW264.7 macrophages (Fig.1D). Upon quantification of the same, we observe that there is a decrease in mean fluorescence intensity with STX7, 6h post-infection on comparison with 2h, followed by an increase at 10h (Fig.1E). Thus, recruitment of STX7 on the SCV exhibits a bimodal pattern, which is in line with its transcript and protein expression profile during infection. We also noticed a significant overlap in the staining pattern of STX7 and LAMP1 in both uninfected and infected cells. Quantification of the same showed that LAMP1 co-localization with STX7 increases at 6h and 10h compared with 2h post-infection (Fig. S1), indicating that they might interact directly or indirectly.

**Figure 1:**
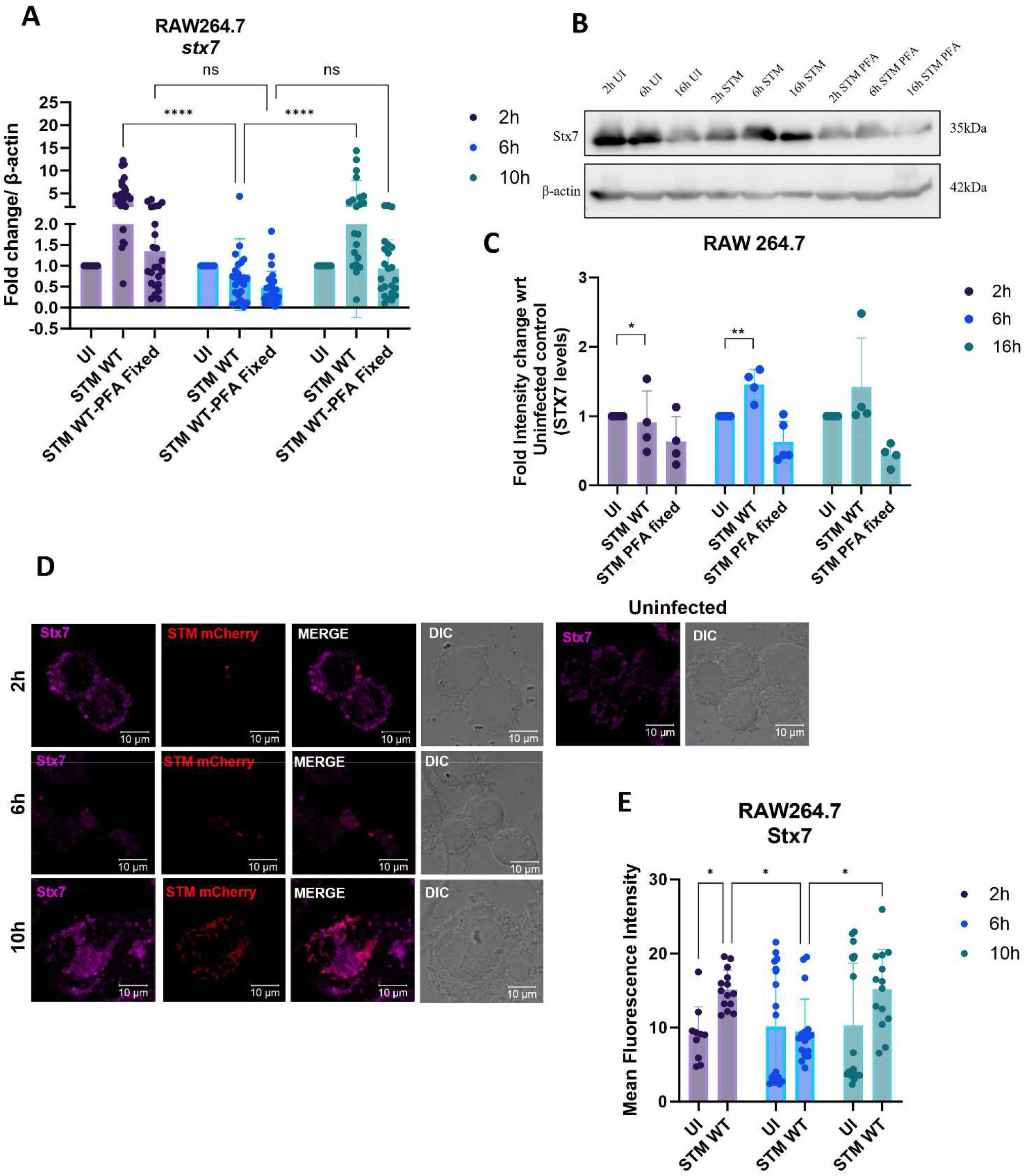
The level of STX7 shows bi-modal regulation during infection with STM WT in macrophages (A) We performed Quantitative RT-PCR to assess the levels of STX7 in infected RAW264.7 macrophages with STM WT or STM WT-PFA-fixed. (B) A representative western blot will be used to assess the levels of STX7 in infected RAW264.7 macrophages with STM WT or STM WT-PFA-fixed. (C) Quantification of (B) with three independent biological replicates. (D) Representative confocal microscopy images of RAW 264.7 cells infected with STM WT and fixed at different time points, stained with STX7 and (E) Quantification of STX7 mean fluorescence intensity (MFI). The scale bar in microscopic images is 10μm, and data is representative of one experiment with more than 50 cells analysed for each condition. All experiments were repeated at least three times, N=3. Student’s unpaired t-test performed for statistical analysis, mean ± SEM/SD * p<0.05, ** p<0.01 and *** p<0.001,

### 2.2 Only the transcript levels of STX7 increase with the progression of STM infection of epithelial cells

The intestinal lining serves as one of the first barriers breached by *Salmonella* during infection, with epithelial cells being key targets for bacterial uptake to access the underlying Peyer’s patches. To explore the role of STX7 in epithelial cells during *Salmonella* infection, we examined its transcript and protein levels in infected HeLa cells. Our analysis revealed an increase in STX7 transcript levels at the late stage of infection (16 h p.i.) with STM WT (Fig. 2A). Interestingly, in cells infected with PFA-fixed STM WT, the transcript levels peaked at an intermediate stage (6 h p.i.) (Fig. 2A). However, STX7 protein levels remained stable throughout the infection and did not display a differential pattern (Fig. 2B-C).

**Figure 2:**
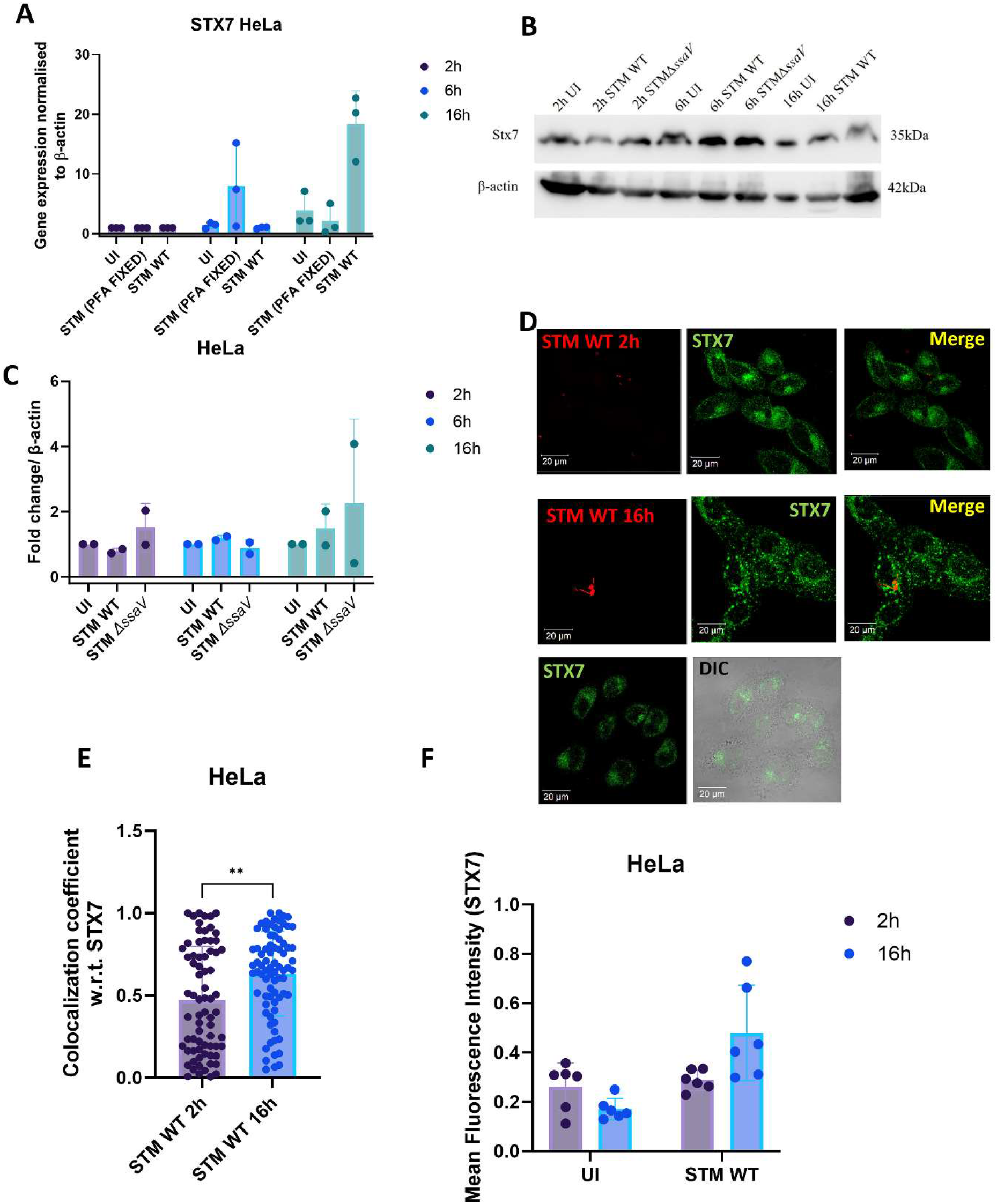
Only the transcript levels of STX7 increase with the progression of STM infection of epithelial cells (A) We performed Quantitative RT-PCR to assess the levels of STX7 in infected HeLa cells with STM WT or STM WT-PFA-fixed. (B) A representative western blot will be used to assess the levels of STX7 in infected HeLa cells with STM WT or STM Δ*ssaV*. (C) Quantification of (B) with three independent biological replicates. (D) Representative confocal microscopy images of HeLa cells infected with STM WT and fixed at different time points, stained with STX7 and (E) Quantification of STX7 co-localization with the bacterium and (F) Quantification of the mean fluorescence intensity (MFI) of STX7. The scale bar in microscopic images is 20μm, and data is representative of one experiment with more than 50 cells analysed for each condition. All experiments were repeated at least three times, N=3. Student’s unpaired t-test performed for statistical analysis, mean ± SEM/SD * p<0.05, ** p<0.01 and *** p<0.001,

Next, we assessed the localization of STX7 in uninfected cells and infected cells at different infection stages in HeLa cells. We observed a significant increase in SCV co-localization with STX7 at the late stage (16 h p.i.) (Fig. 2D-E), despite no significant changes in STX7 levels as measured by mean fluorescence intensity (Fig. 2F). This marked shift in localization suggests a functional reorganization of STX7 during infection. Together, these findings highlight a potential role for STX7 in epithelial cells during *Salmonella* infection, particularly in SCV biogenesis or maintenance.

### 2.3 Knockdown and overexpression of STX7 impact *Salmonella* proliferation and phagocytosis in macrophages and epithelial cells

To investigate the role of STX7 in *Salmonella* pathogenesis, we performed knockdown experiments using two independent shRNAs -shRNA1 and shRNA2 plus a mix of shRNA1 and shRNA2 and overexpression studies with STX7-FL (full-length)[27], STX7-cyto (membrane anchoring domain deleted therefore protein remain cytosolic act as a negative control[24]), and an empty vector (eGFP-EV) control. The experiments were conducted in macrophages (RAW264.7 cells) and epithelial cells (HeLa cells) to assess the effects on both bacterial proliferation and phagocytosis. STX7 knockdown efficiency was validated by immunofluorescence analysis and qPCR and in both cell types (Fig S2-S3). Following knockdown with shRNA1 and shRNA2, phagocytosis assays revealed a significant decrease in bacterial uptake/invasion in STX7-depleted cells. RAW264.7 macrophages exhibited reduced bacterial internalization with both shRNA constructs compared to control cells (Fig. 3A). A similar trend was observed in HeLa cells as percentage invasion (Fig 3E), where STX7 knockdown led to a marked decline in *Salmonella* uptake. This indicates that STX7 may facilitate bacterial entry into host cells.

**Figure 3:**
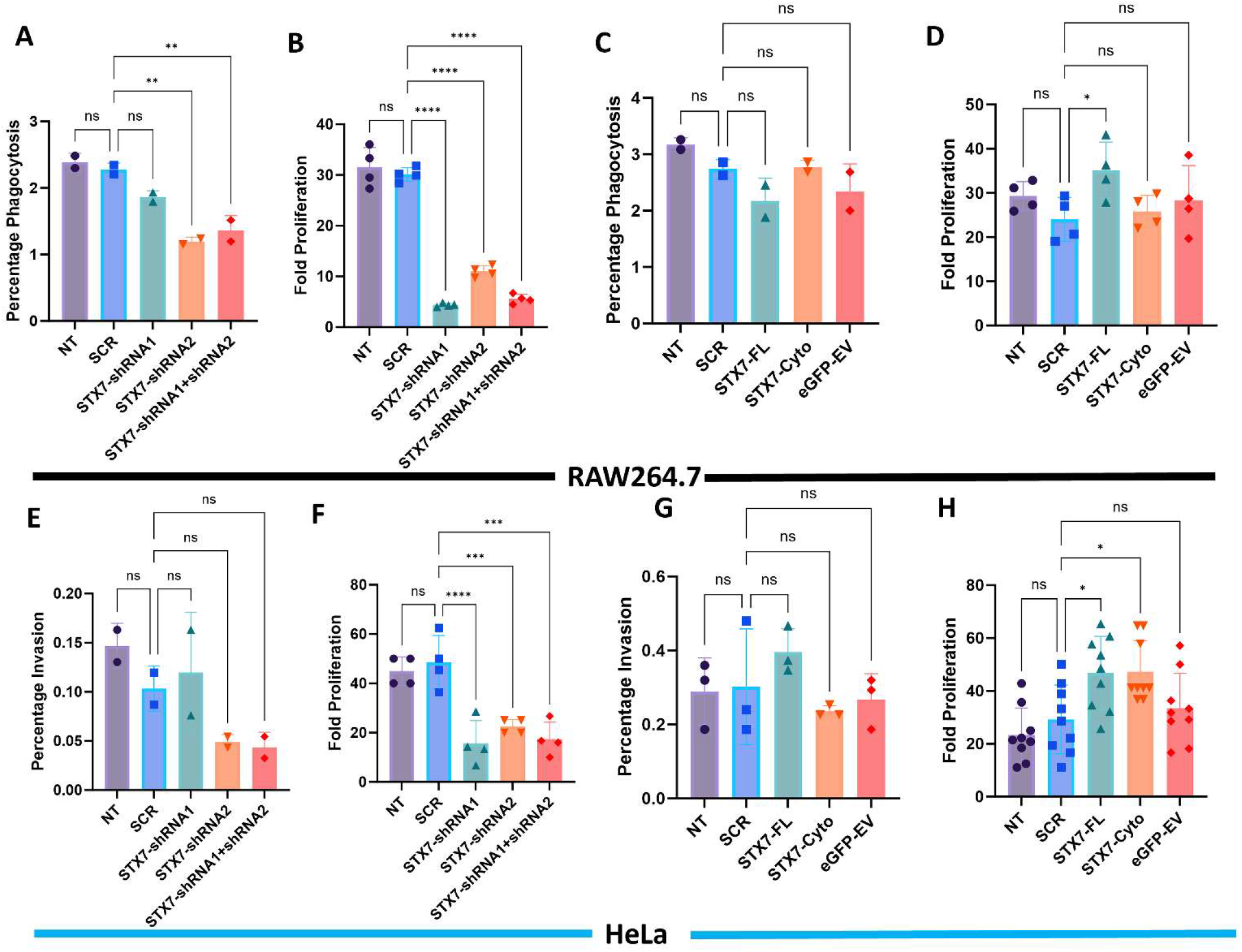
Knockdown and overexpression of STX impact *Salmonella* proliferation and phagocytosis in macrophages and epithelial cells We performed an intracellular survival assay in STX7 knockdown RAW264.7 macrophages (using either shRNA1 or shRNA2 or a 1:1 mix of shRNA1 and shRNA2) (A) percentage phagocytosis (B) fold proliferation. We also overexpressed the full-length STX7 (STX7-FL), or cytoplasmic STX7 (truncated negative control) or eGFP- empty vector control (eGFP-EV) in RAW264.7 macrophages (C) percentage phagocytosis and (D) fold proliferation. Similarly, we also performed an intracellular survival assay in STX7 knockdown HeLa cells (using either shRNA1 or shRNA2 or a 1:1 mix of shRNA1 and shRNA2) (A) percentage phagocytosis (B) fold proliferation. We also overexpressed the full-length STX7 (STX7-FL), or cytoplasmic STX7 (truncated negative control) or eGFP- empty vector control (eGFP-EV) in HeLa cells to assess the (C) percentage phagocytosis and (D) fold proliferation. All experiments were repeated at least three times, N=3. Student’s unpaired t-test performed for statistical analysis, mean ± SEM/SD, * p<0.05, ** p<0.01 and *** p<0.001,

A significant reduction in *Salmonella* fold proliferation was observed in RAW264.7 macrophages (Fig. 3B) and HeLa epithelial cells compared to controls (non-targeting shRNA) (Fig. 3F). In macrophages, the fold proliferation of *Salmonella* is significantly decreased by shRNA1, shRNA2, and a mix of shRNA1 and shRNA2, respectively. Similarly, in HeLa cells, there was a significant decrease, too. This suggests that STX7 is critical for *Salmonella’s* intracellular survival and replication, too.

Overexpression of STX7-FL in both cell types resulted in increased *Salmonella* proliferation and phagocytosis compared to the eGFP-EV control (Fig 3C-D, G-H). In contrast, overexpression of STX7-cyto did not show any significant changes, indicating that the full-length STX7 membrane-bound protein, is required for *Salmonella* survival and uptake (Fig. 3D-E). This highlights the functional importance of the intact STX7 protein.

While STX7 knockdown and overexpression showed a consistent trend across both cell types, the magnitude of changes differed. RAW264.7 macrophages demonstrated a more pronounced impact on bacterial proliferation compared to HeLa cells (Fig. 3), possibly reflecting cell-type-specific differences in STX7-associated pathways or *Salmonella* survival strategies.

### 2.4. Expression of STX7 Adaptor/Binding Partners and Its Association with the SCV During Infection

To further elucidate the role of STX7 in *Salmonella* pathogenesis, we assessed the expression profiles of key STX7 adaptors and binding partners during infection. Additionally, live-cell imaging experiments were conducted to examine the dynamics of STX7 association with the *Salmonella*-containing vacuole (SCV) at various stages of infection. Snapshots from live-cell microscopy (Videos 1 and 2) illustrate the interaction of STX7 with the SCV at early (2h post-infection [p.i.]) and late (16h p.i.) stages of infection.

The expression levels of STX7 adaptor/binding partners were quantified at different stages of infection in RAW264.7 cells (Fig. 4A-D). At the early stage (2 h p.i.), there was a modest increase in the transcript levels of these adaptors compared to uninfected controls. The expression levels peaked at the intermediate stage (6 h p.i.) before gradually decreasing by the late stage (16 hp.i.). This temporal pattern suggests a dynamic regulation of STX7-binding partners during infection, which may reflect their functional roles in facilitating *Salmonella* survival and replication within the SCV.

**Figure 4:**
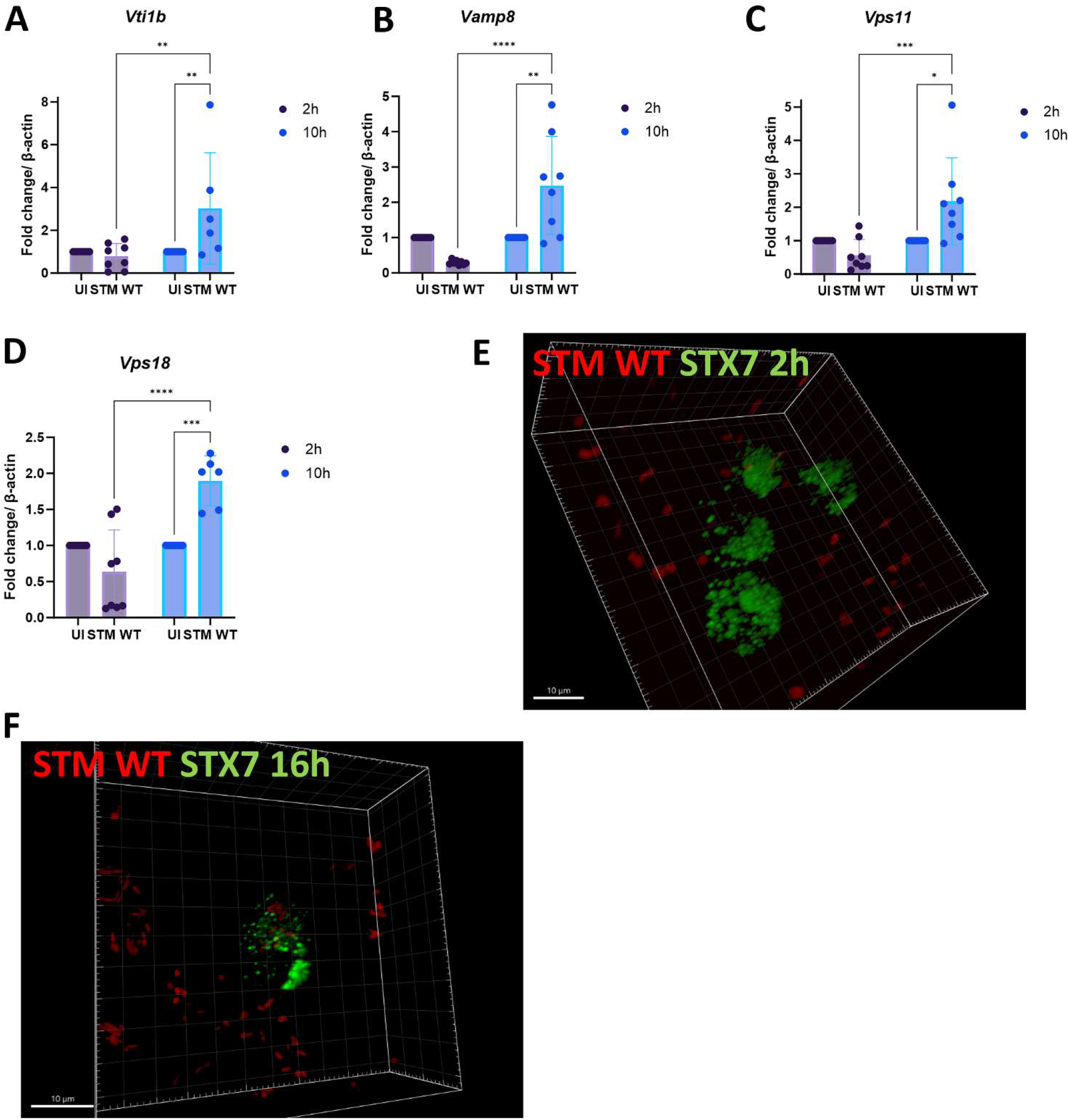
Expression of STX7 Adaptor/Binding Partners and Its Association with the SCV During Infection (A-D) Quantitative RT-PCR analysis showing fold change in mRNA expression levels of Vti1b (A), Vamp8 (B), Vps11 (C), and Vps18 (D) normalized to β-actin in unstimulated (UI) and Salmonella-infected (STM WT) macrophages at 2 hours and 10 hours post-infection. All experiments were performed in biological triplicates (N=3), and the data are represented as mean ± SEM. Statistical analysis was conducted using a Student’s unpaired t-test* p<0.05, ** p<0.01 and *** p<0.001. (E-F) 3D live-cell imaging snapshots of macrophages infected with Salmonella WT (red) showing STX7 (green) localization. (E) At 2 hours post-infection, STX7 co-localizes with Salmonella in discrete vesicular structures. (F) At 16 hours post-infection, the majority of STX7-positive vesicles have aggregated, indicating enhanced recruitment. Scale bar = 10 μm.

To visualize the association of STX7 with the SCV, we performed live-cell microscopy using HeLa cells expressing full-length STX7 (STX7-FL) fused to a fluorescent tag. Videos 1 and 2 provide real-time visualization of the interaction between STX7 and the SCV during infection, and representative snapshots are included in Fig. 4E and 4F.

At 2 h p.i. (early stage), we observed a strong co-localization of STX7 with the SCV, indicating that STX7 is rapidly recruited to the site of infection (Fig. 4E). This association persisted and became more pronounced at the late stage (16 h p.i.) (Fig. 4F). These findings suggest that STX7 may play an active role in modulating SCV dynamics, particularly at early and late stages of infection. Quantitative analysis of live-cell microscopy data revealed a significant increase in the co-localization of STX7 with the SCV at 2h and 16 h p.i. compared to uninfected cells (Fig. 4E-F). This was accompanied by a consistent intensity of STX7 fluorescence within the SCV, indicating stable recruitment of STX7 over time. These results highlight the involvement of STX7 in SCV maturation and its potential role in facilitating *Salmonella* survival within host epithelial cells.

The expression analysis and live-cell microscopy collectively demonstrate that STX7 and its adaptors are dynamically regulated during *Salmonella* infection. The strong and sustained association of STX7 with the SCV, particularly at early and late stages of infection, suggests that STX7 may contribute to SCV stabilization or maturation. These data provide further evidence for the critical role of STX7 in the intracellular life cycle of *Salmonella*.

## 3. Discussion

Our findings reveal a crucial role for STX7 in modulating *Salmonella* infection dynamics in both macrophages and epithelial cells. Syntaxin 7, a Qa-SNARE protein, is predominantly localized on late endosomes and lysosomes, where it mediates endo-lysosomal fusion and trafficking processes [11]. However, intracellular pathogens, including *Salmonella*, exploit such host cell machinery to ensure their survival and replication within host cells [1]. Previous reports have highlighted the ability of *Salmonella* to hijack host SNARE proteins, such as STX3 and STX4, to modulate vacuole maturation and nutrient acquisition [5–7, 9] Our study extends these findings by establishing STX7 as a critical player in the intracellular pathogenesis of *Salmonella* Typhimurium.

Our data demonstrate a bimodal regulation of STX7 at both the transcript and protein levels during *Salmonella* infection in macrophages, with peaks observed at 2h and 10h post-infection and a transient decline at 6h. This dynamic regulation might reflect distinct stages of SCV biogenesis and maturation, as *Salmonella* actively remodels the SCV to prevent lysosomal fusion while acquiring nutrients [4]. Immunofluorescence studies confirmed that STX7 is recruited to the SCV during all stages of infection, with significant co-localization observed with LAMP1, a late endosome and lysosome marker. The increased LAMP1 co-localization at 6h and 10h suggests a potential interaction between STX7 and late endosomal/lysosomal compartments, which *Salmonella* may exploit to modulate vesicle trafficking or nutrient acquisition.

Interestingly, STX7 transcript levels in epithelial cells increased significantly during late-stage infection (16h post-infection), but protein levels remained stable throughout. Despite this, the co-localization of STX7 with SCVs increased at the late stage, indicating a functional reorganization of STX7 without significant changes in its expression. This discrepancy between transcriptional and localization dynamics might reflect cell type-specific regulatory mechanisms or differences in SCV composition between macrophages and epithelial cells.

Our knockdown and overexpression experiments underscore the functional importance of STX7 in *Salmonella* infection. Knockdown of STX7 significantly reduced bacterial invasion and intracellular proliferation in both macrophages and epithelial cells, indicating its critical role in bacterial entry and intracellular survival. Conversely, overexpression of full-length STX7 enhanced both *Salmonella* proliferation and phagocytosis, further supporting the notion that *Salmonella* hijacks STX7 to establish its intracellular niche. Notably, overexpression of a truncated, cytoplasmic version of STX7 did not produce similar effects, emphasizing the importance of the intact protein and its subcellular localization for *Salmonella* pathogenesis.

The ability of *Salmonella* to manipulate STX7 aligns with previous reports of intracellular pathogens exploiting host endocytic pathways. For example, *Legionella* cleaves STX17 to block autophagy [2], and *Chlamydia* employs SNARE-like proteins to modulate host vesicle trafficking [3]. Our study highlights that *Salmonella* follows a similar strategy by exploiting STX7 to manipulate the SCV and evade lysosomal degradation.

## 4. Conclusion

In summary, our study provides compelling evidence that STX7 plays a critical role in *Salmonella* pathogenesis by facilitating bacterial entry, SCV biogenesis, and intracellular survival. These findings deepen our understanding of the intricate host-pathogen interactions that underpin *Salmonella* infection and highlight STX7 as a promising target for therapeutic intervention.

## 5. Material and Methods-

### 5.1 Bacterial strains and growth condition

*Salmonella* enterica serovar Typhimurium 14028S [STM (WT)] or STM (WT) constitutively expressing mCherry through pFPV25.1 were used in all experiments. All the bacterial strains were cultured in Luria Bertani (LB) broth (HiMedia) with constant shaking (160 rpm) at 37°C. Ampicillin (50μg/ml) was used wherever required.

### 5.2 Cell culture protocol: DMEM-

Dulbecco’s Modified Eagle Medium (Lonza) supplemented with 10% FBS (Gibco) was used to culture RAW 264.7 murine macrophages and HeLa cells. The cells were maintained at 37°C with 5% CO2, in a humidified incubator. Cells were seeded into the wells of 24 (for confocal microscopy and intracellular survival assay) or 6 well plate (for RNA isolation) as per requirement at a confluency of 60-70%, preceding the experiment.

### 5.3 Gentamicin protection assay

Macrophages were infected with stationary-phase bacterial culture with MOI of 10 or 25 (for confocal study). For synchronisation of the infection and enhancing bacterial adhesion, tissue culture plates were subjected to centrifugation at 600rpm for 5 minutes, followed by incubation for 25 minutes at 37°C in a humidified incubator with 5% CO2. Post incubation cells were washed with PBS and fresh media (DMEM + 10% FBS) containing 100μg/ml gentamicin was added, followed by 1h of incubation at 37°C and 5% CO2. Subsequently, the gentamicin concentration was reduced to 25μg/ml and maintained until the cells were harvested.

### 5.4 RNA isolation and quantitative RT PCR

Total RNA was isolated at specific time points post infection by using TRIzol (RNAiso plus Takara) as per manufacturer’s protocol. Quantification of RNA was performed in Nano Drop (Thermo-Fischer scientific). The quality of isolated RNA was detected by performing gel electrophoresis with 2% agarose gel. 2μg of RNA was subjected to DNase treatment at 37°C for 1 hr followed by addition of EDTA and heat inactivation at 75°C for 10 mins. The mRNA was reverse transcribed to cDNA using oligo dT primer, buffer, dNTPs and reverse transcriptase (Takara) as per manufacturer’s protocol. The expression profile of target genes was evaluated using specific primers by using TB green RT-qPCR master mix (Takara) in BioRad Real time PCR instrument or Applied Biosystem QuantStudio-5 system. β-actin was used as an internal control. All the reactions were setup in 384 well plate with two or three technical replicates for each sample.

**Table 1:**
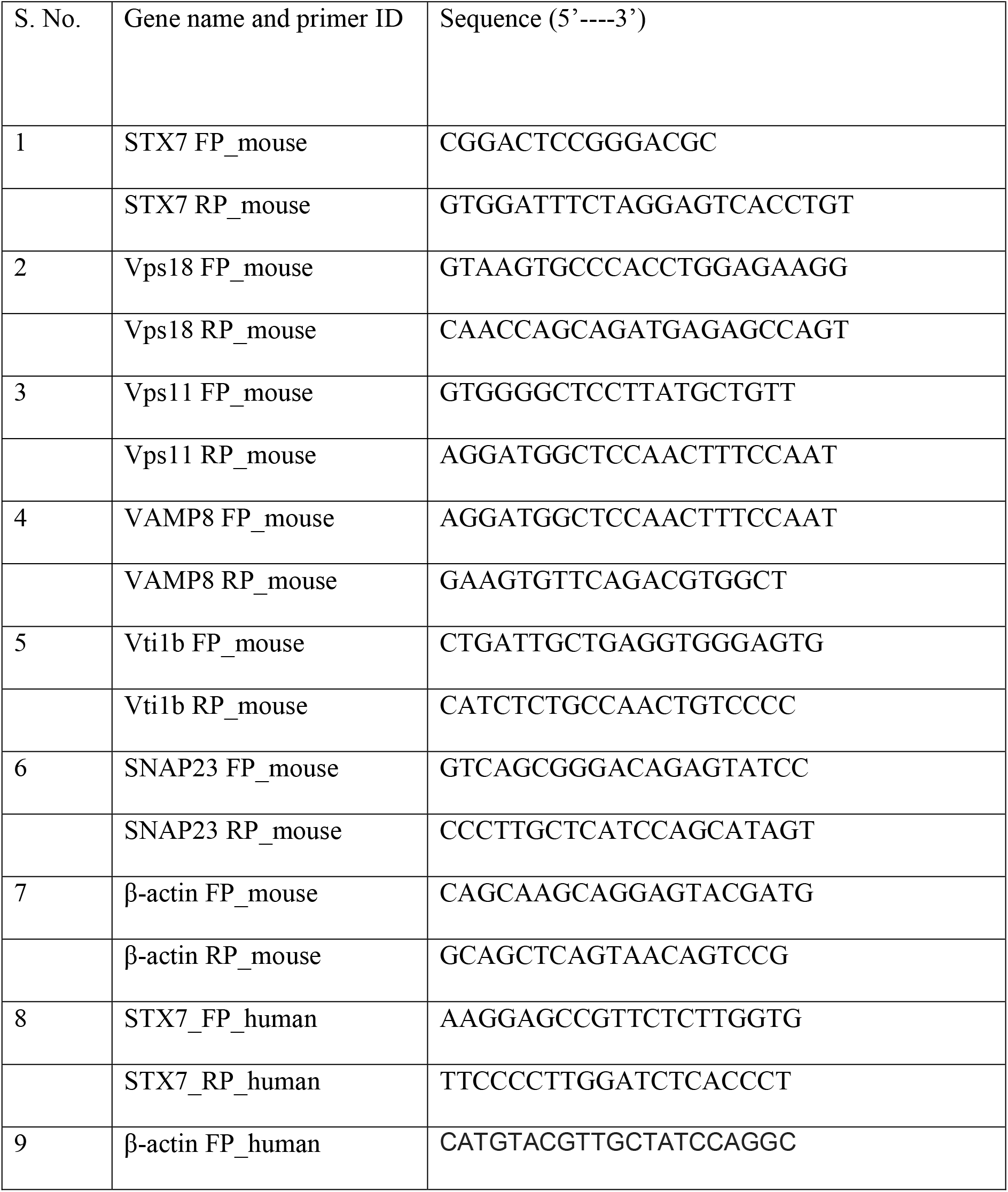
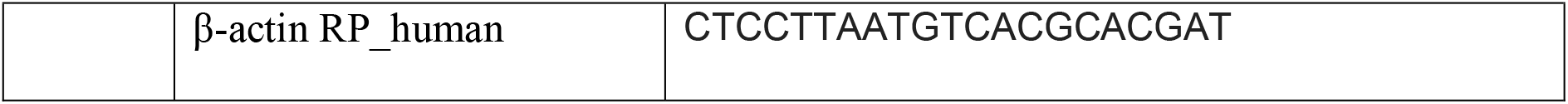
List of Primers

### 5.5 Confocal Microscopy

HeLa or RAW 264.7 cells were seeded in 24 well plates containing coverslips. At the specified time points post-infection with mCherry-tagged STM, cells were washed with PBS and fixed with 3.5% paraformaldehyde for 15 minutes at room temperature. The cells were washed twice with PBS and were incubated with in blocking solution [0.01% saponin (Sigma), 2.5% BSA in PBS] containing primary antibodies against STX7 and LAMP1 overnight at 4°C. Following this, cells were washed with PBS stained with appropriate secondary antibodies tagged with fluorophores for 1 hr at room temperature (RT). The coverslips were then mounted onto a clean glass slide and imaged under confocal microscope (Zeiss LSM-880) using 63X objective. The images were analysed using ZEN black 2012 software.

**Table 2:** List of antibodies

| Sl. No. | Antibody | Catalogue Number |
| --- | --- | --- |
| 1 | Syntaxin 7 Monoclonal Antibody | sc-514017 (SCBT) |
| 2 | Beta-actin-HRP tagged | D6A8 (CST) |
| 3 | Mouse LAMP1 Monoclonal Antibody | 1D4B (DSHB) |

### 5.6 Transfection protocol

Transfection for shRNA-mediated knockdown was carried out by Polyethylenimine (PEI) (Sigma). Plasmids harboring shRNA in panko.2 vector backbones specific to STX7 were used for transfection. Plasmid harbouring scrambled sequence of shRNA, served as a control, was also used for transfection. Plasmid DNA was used at a concentration of 500ng per well in a 24- well plate. Plasmid and PEI were added in 1:2 ratio in serum free DMEM media and incubated for 20 minutes at room temperature. Post incubation, the DNA: PEI cocktail was added to the seeded HeLa or RAW 264.7 macrophages. After 6-8h of incubation, serum-free media was replaced with complete media containing 10% FBS. Post 48h of transfection, transfected cells were either harvested for knockdown confirmation studies or subjected to infection with STM.

**Table 3:**
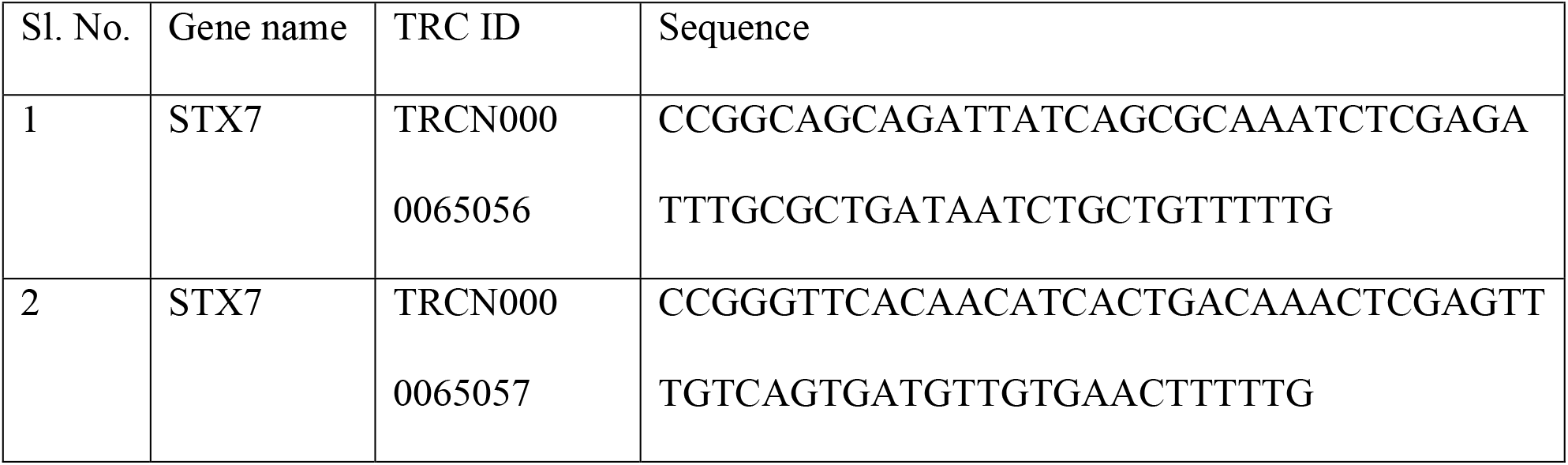
List of shRNA

### 5.7 Percentage phagocytosis and Intracellular proliferation

Following infection of the transfected cells with STM at a MOI of 10, cells were treated with DMEM (Sigma) supplemented with 10% FBS (Gibco) containing 100μg/ml gentamicin for 1h. Subsequently, the gentamicin concentration was reduced to 25μg/ml and maintained until the specified time point. 2h, 6h, 10h, and 16h post-infection, cells were lysed in 0.1% triton-X-100. Lysed cells and the pre-inoculum were serially diluted and plated onto *Salmonella*- Shigella agar to obtain colony-forming units (CFU). Calculations for percentage phagocytosis and fold proliferation are as follows:

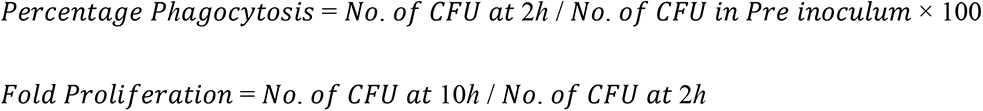

### 5.8 Statistical analysis

Data was analysed and graphed using the GraphPad Prism 8 software (San Diego, CA). Statistical significance was determined by Student’s t-test to obtain p values. Adjusted p-values below 0.05 are considered statistically significant. The results are expressed as mean ± SEM, unless mentioned otherwise.

## Acknowledgements and funding

We would like to thank Professor Sunando Datta, IISER Bhopal, India, for kindly providing pEGFPN1-STX7 (full length) construct and Professor Reinhard Jahn, Max Planck Institute for Biophysical Chemistry, Göttingen, Germany, for kindly permitting to use pEGFPN1- STX7cyto construct (the pEGFPN1-STX7cyto plasmid was provided us by Prof. Datta with due permission from Prof. Jahn). Divisional and Departmental Confocal Facility and Departmental Real-Time PCR Facility at MCB, IISc are duly acknowledged. Mr. Sumeeth, Mrs. Saima, and Ms. Navya are acknowledged for their help with image acquisition. This work was supported by the Department of Biotechnology (DBT), Ministry of Science and Technology, the Department of Science and Technology (DST), Ministry of Science and Technology. The authors jointly acknowledge the DBT-IISc partnership program. Infrastructure support from ICMR (Center for Advanced Study in Molecular Medicine), DST (FIST), UGC-CAS (special assistance), and TATA fellowship is acknowledged. This work was also supported by the Department of Biotechnology (BT/PR32489/BRB/10/1786/2019), Science and Engineering Research Board (CRG/2019/000281), DBT-NBACD (BT/HRD-NBA-NWB/38/2019-20) and India Alliance (500122/Z/09/Z) to SRGS. RV, RC and DH duly acknowledge the CSIR-SRF fellowship. AVN acknowledges the IISc-MHRD fellowship, and AS acknowledges the UGC-SRF fellowship.

## References

1. Chatterjee, R., S.R.G. Setty, and D. Chakravortty, SNAREs: a double-edged sword for intravacuolar bacterial pathogens within host cells. Trends Microbiol, 2024. 32(5): p. 477–493.

2. Arasaki, K., et al., Legionella effector Lpg1137 shuts down ER-mitochondria communication through cleavage of syntaxin 17. Nat Commun, 2017. 8: p. 15406.

3. Delevoye, C., et al., SNARE protein mimicry by an intracellular bacterium. PLoS Pathog, 2008. 4(3): p. e1000022.

4. Knuff, K. and B.B. Finlay, What the SIF Is Happening-The Role of Intracellular Salmonella-Induced Filaments. Front Cell Infect Microbiol, 2017. 7: p. 335.

5. Madan, R., et al., Salmonella acquires lysosome-associated membrane protein 1 (LAMP1) on phagosomes from Golgi via SipC protein-mediated recruitment of host Syntaxin6. J Biol Chem, 2012. 287(8): p. 5574–87.

6. Singh, P.K., et al., Salmonella SipA mimics a cognate SNARE for host Syntaxin8 to promote fusion with early endosomes. J Cell Biol, 2018. 217(12): p. 4199–4214.

7. Chatterjee, R., et al., Syntaxin 3 SPI-2 dependent crosstalk facilitates the division of Salmonella containing vacuole. Traffic, 2023. 24(7): p. 270–283.

8. Chatterjee, R., et al., Deceiving The Big Eaters: Salmonella Typhimurium SopB subverts host cell Xenophagy in macrophages via dual mechanisms. Microbes Infect, 2023: p. 105128.

9. Stevenin, V., et al., Dynamic Growth and Shrinkage of the Salmonella-Containing Vacuole Determines the Intracellular Pathogen Niche. Cell Rep, 2019. 29(12): p. 3958–3973 e7.

10. Budak, G., et al., Reconstruction of the temporal signaling network in Salmonella-infected human cells. Front Microbiol, 2015. 6: p. 730.

11. Mullock, B.M., et al., Syntaxin 7 is localized to late endosome compartments, associates with Vamp 8, and Is required for late endosome-lysosome fusion. Mol Biol Cell, 2000. 11(9): p. 3137–53.

12. D’Costa, V.M., et al., BioID screen of Salmonella type 3 secreted effectors reveals host factors involved in vacuole positioning and stability during infection. Nat Microbiol, 2019. 4(12): p. 2511–2522.

13. Kasai, K. and K. Akagawa, Roles of the cytoplasmic and transmembrane domains of syntaxins in intracellular localization and trafficking. J Cell Sci, 2001. 114(Pt 17): p. 3115–24.

14. Prekeris, R., et al., Differential roles of syntaxin 7 and syntaxin 8 in endosomal trafficking. Mol Biol Cell, 1999. 10(11): p. 3891–908.

15. Pryor, P.R., et al., Combinatorial SNARE complexes with VAMP7 or VAMP8 define different late endocytic fusion events. EMBO Rep, 2004. 5(6): p. 590–5.

16. Dingjan, I., et al., Oxidized phagosomal NOX2 complex is replenished from lysosomes. J Cell Sci, 2017. 130(7): p. 1285–1298.

17. Chidambaram, S., J. Zimmermann, and G.F. von Mollard, ENTH domain proteins are cargo adaptors for multiple SNARE proteins at the TGN endosome. J Cell Sci, 2008. 121(Pt 3): p. 329–38.

18. Achuthan, A., et al., Regulation of the endosomal SNARE protein syntaxin 7 by colony- stimulating factor 1 in macrophages. Mol Cell Biol, 2008. 28(20): p. 6149–59.

19. Pirooz, S.D., et al., UVRAG is required for virus entry through combinatorial interaction with the class C-Vps complex and SNAREs. Proc Natl Acad Sci U S A, 2014. 111(7): p. 2716–21.

20. Suzuki, J., et al., Involvement of syntaxin 7 in human gastric epithelial cell vacuolation induced by the Helicobacter pylori-produced cytotoxin VacA. J Biol Chem, 2003. 278(28): p. 25585–90.

21. Kehl, A., et al., A trafficome-wide RNAi screen reveals deployment of early and late secretory host proteins and the entire late endo-/lysosomal vesicle fusion machinery by intracellular Salmonella. PLoS Pathog, 2020. 16(7): p. e1008220.

22. Pha, K., et al., The Chlamydia effector IncE employs two short linear motifs to reprogram host vesicle trafficking. Cell Rep, 2024. 43(8): p. 114624.

23. Chatterjee, R., et al., From Eberthella typhi to Salmonella Typhi: The Fascinating Journey of the Virulence and Pathogenicity of Salmonella Typhi. ACS Omega, 2023. 8(29): p. 25674–25697.

24. Antonin, W., et al., A SNARE complex mediating fusion of late endosomes defines conserved properties of SNARE structure and function. EMBO J, 2000. 19(23): p. 6453–64.

25. Brumell, J.H., et al., Disruption of the Salmonella-containing vacuole leads to increased replication of Salmonella enterica serovar typhimurium in the cytosol of epithelial cells. Infect Immun, 2002. 70(6): p. 3264–70.

26. Roark, E.A. and K. Haldar, Effects of lysosomal membrane protein depletion on the Salmonella-containing vacuole. PLoS One, 2008. 3(10): p. e3538.

27. Parveen, S., et al., Syntaxin 7 contributes to breast cancer cell invasion by promoting invadopodia formation. J Cell Sci, 2022. 135(12).

